# Interactions Between the Frequency of the Duffy Antigen and the Dynamics of *P. vivax* Malaria Infections

**DOI:** 10.1101/2025.11.30.691431

**Authors:** Elizabeth Ghartey, Gautam Rai, Dasha Selivonenko, Rachel K. Wissenbach, Joan Ponce

**Affiliations:** University of Arizona, Tucson AZ, United States; Arizona State University, Tempe AZ, United States; Rice University, Houston TX, United States

## Abstract

The malarial parasite *Plasmodium vivax* has infected humans for millennia. As such, alleles like the FY*O allele in the Duffy Antigen Receptor for Chemokines (DARC) and the sickle cell allele (HbS) have been naturally selected for in malaria-endemic regions because they confer resistance to malaria, thereby increasing the survival and reproductive success of individuals carrying these protective alleles. As resistance becomes more common in a population, malaria incidence is expected to decline, reducing the evolutionary pressure for additional resistance. In this work, we explore the interaction between these processes. We construct a model with seasonality that tracks the frequency of Duffy genotypes and leverages fast/slow dynamics to analyze the coupled dynamics of malaria transmission and changes in the gene frequency of the DARC genotype. Specifically, we investigate how the burden of malaria changes with the fractions of people with the various DARC genotypes. We derive the basic reproduction number as the threshold condition for the stability of the disease-free equilibrium and interpret the *R*_0_ as a weighted sum of the cases generated by infected individuals of each genotype. Additionally, we calibrate our model using data from the Amazonas region in Brazil, which has a polymorphic population with respect to DARC, and still reports a substantial number of *P. vivax* cases. Analysis of our model determines the proportion of the population that must be Duffy-negative in order for the entire population to be protected against *P. vivax* without any further interventions. Furthermore, we assess how different proportions of Duffy-negative individuals influence the monthly incidence of *P. vivax* cases.

## 1 Introduction

### 1.1 Influence of Genetics on the Epidemiology of Malaria

Malaria is an infectious disease spread by mosquitoes that are infected with *Plasmodium* parasites. Whereas *P. falciparum* is the most dangerous strain and the most prevalent in Africa, *P. vivax* is the most geographically widespread strain of malaria and the most common in the rest of the world [12]. Malaria is a perennial affliction to many tropical countries: in 2022, it caused an estimated 249 million cases globally, resulting in more than six hundred thousand deaths and billions of dollars in global costs [22].

The selective pressure induced by malaria has demonstrably influenced the genetic composition of tropical populations. Examples include hemoglobinopathies like the sickle cell trait in Sub-Saharan Africa and *β*-thalassemia in the Eastern Mediterranean. The sickle cell trait is caused by a missense mutation of the *β*-globin gene, resulting in the abnormal synthesis of the *β*-globin protein. The vast majority of people with sickle cell trait and sickle cell disease live in Sub-Saharan Africa. Similarly, *β*-thalassemia is characterized by the obstructed synthesis of the *β*-globin chain, which causes numerous phenotypes that may be asymptomatic or debilitating. People with sickle cell trait and some phenotypes of *β*thalassemia have significant protection against *P. falciparum* malaria compared to people without either condition [5, 7, 10].

Another example of an adaptation against malaria is the absence of the Duffy antigen receptor for chemokines (DARC) on human red blood cells. *P. vivax* protozoa rely on the interaction between DARC and the *Plasmodium vivax* Duffy-binding protein (PvDBP) to invade human reticulocytes. Having the Duffy-negative trait thereby removes a mode of invasion into reticulocytes by *P. vivax* protozoa. Nonetheless, infections by *P. vivax* have been observed in Duffy-negative individuals, but the mechanism of infection in these cases are not fully understood [19].

The transmission patterns of malaria are highly influenced by the Duffy-negative trait. Duffy-negativity is very common in sub-Saharan Africa, so the fraction of the population at risk of *vivax* malaria is very small. Consequently, the majority of malaria infections are caused by *P. falciparum*. Although *P. vivax* infections is exceedingly rare in sub-Saharan Africa, it is the most common malarial parasite in the rest of the world [9].

Our system models the dynamics of *P. vivax* malaria in the Amazonas state of Brazil, because this population is much more polymorphic with respect to the Duffy antigen trait than most parts of the world. Kano et al. [15] conducted a longitudinal study on the influence of DARC on people’s susceptibility to *vivax* malaria in Rio Pardo, which is located in the Brazilian state of Amazonas. In this study, they sampled antibody response to *vivax* malaria, and disaggregated it by Duffy genotype to ascertain the influence of natural immunity. We used data from the longitudinal work of Kano et al to determine the populations of interest in our study.

In this paper, we use ordinary differential equations to characterize the dynamics of *vivax* malaria in Rio Pardo, Brazil, with relation to the Duffy antigen. We reference previous population genetics models that relate to the sickle cell trait. Feng et al. [6] used a system of ordinary differential equations to describe the relationship between malaria transmission dynamics and the genetic frequency of the S-gene that causes sickle cell anemia. Furthermore, Feng and Castillo-Chavez [5] used an expanded system of ordinary differential equations with three elements to characterize the complex dynamics of the S-gene beyond the standard two-element system.

We use the Rio Pardo area studied by Kano et al. [15], because it is more polymorphic than most parts of the world. This has implications for transmission dynamics of *vivax* malaria in the region. Figure 1 denotes the genotypic frequencies of the Duffy trait in Rio Pardo.

**Figure 1.**
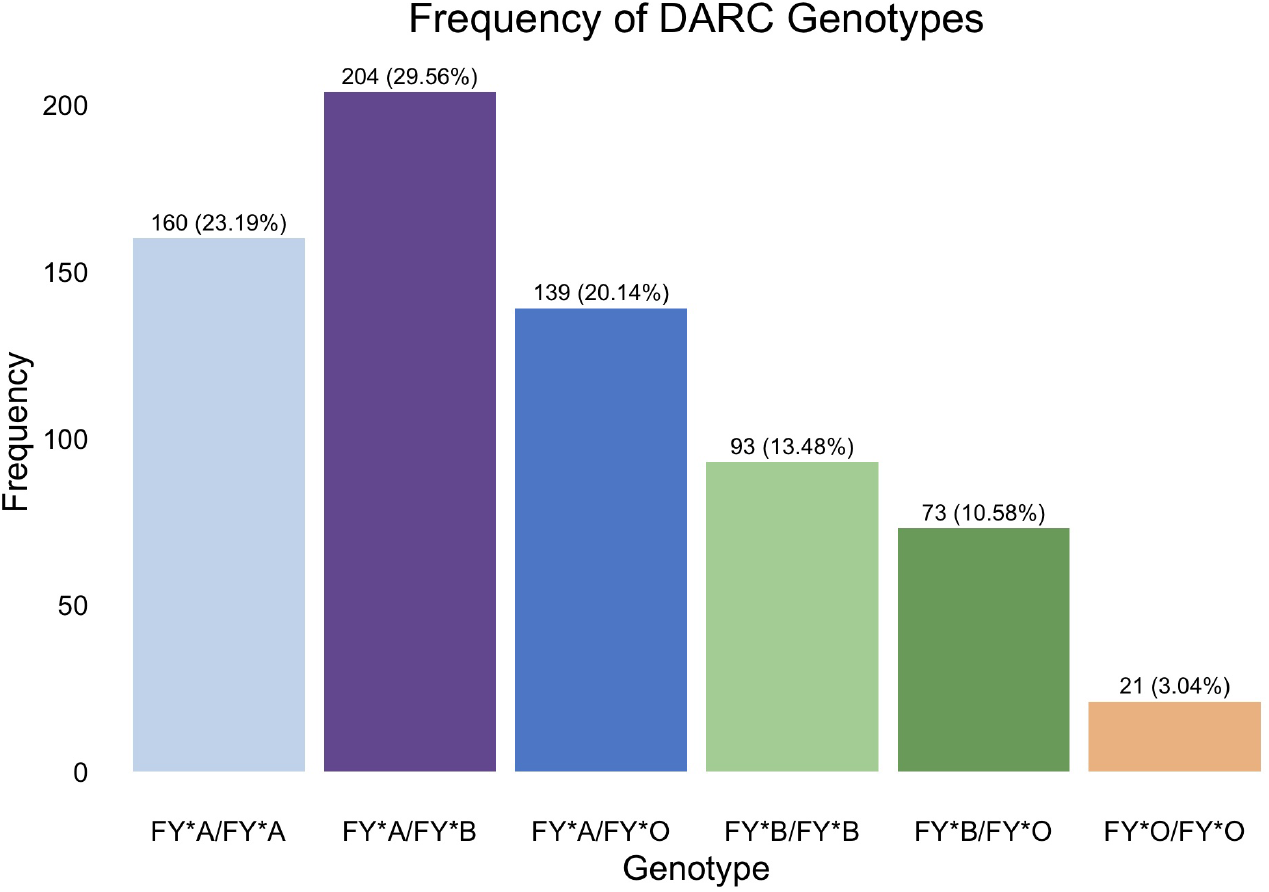
Distribution of Duffy antigen polymorphisms in Rio Pardo, Amazonas, Brazil.

The above graph shows that the large majority of people in Rio Pardo are Duffy-positive, but a considerable minority are also homozygous Duffy-negative. The characteristics of the Duffy antigen suggest that people who are Duffy negative (FY*O/FY*O) are highly unlikely to contract *vivax* malaria. This is supported by Figure 2, which shows the *vivax* malaria cases by genotype in the Kano study [15].

**Figure 2.**
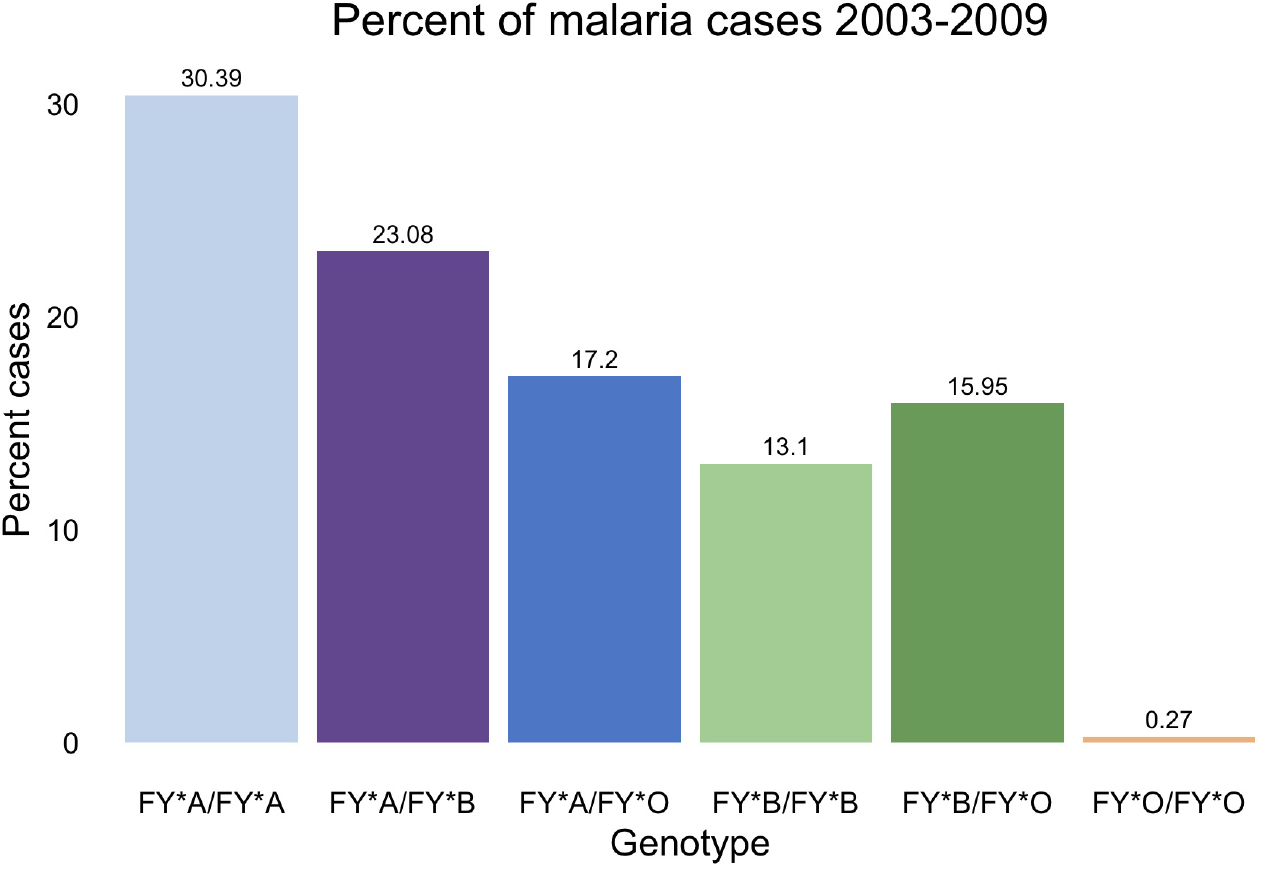
Malaria cases by genotype in a longitudinal study conducted in Rio Pardo, Amazonas, Brazil. Besides corroborating the protective effects of the Duffy-negative trait, this data also suggests that the FY*B trait has limited protective effects against *vivax* malaria.

**Figure 3.**
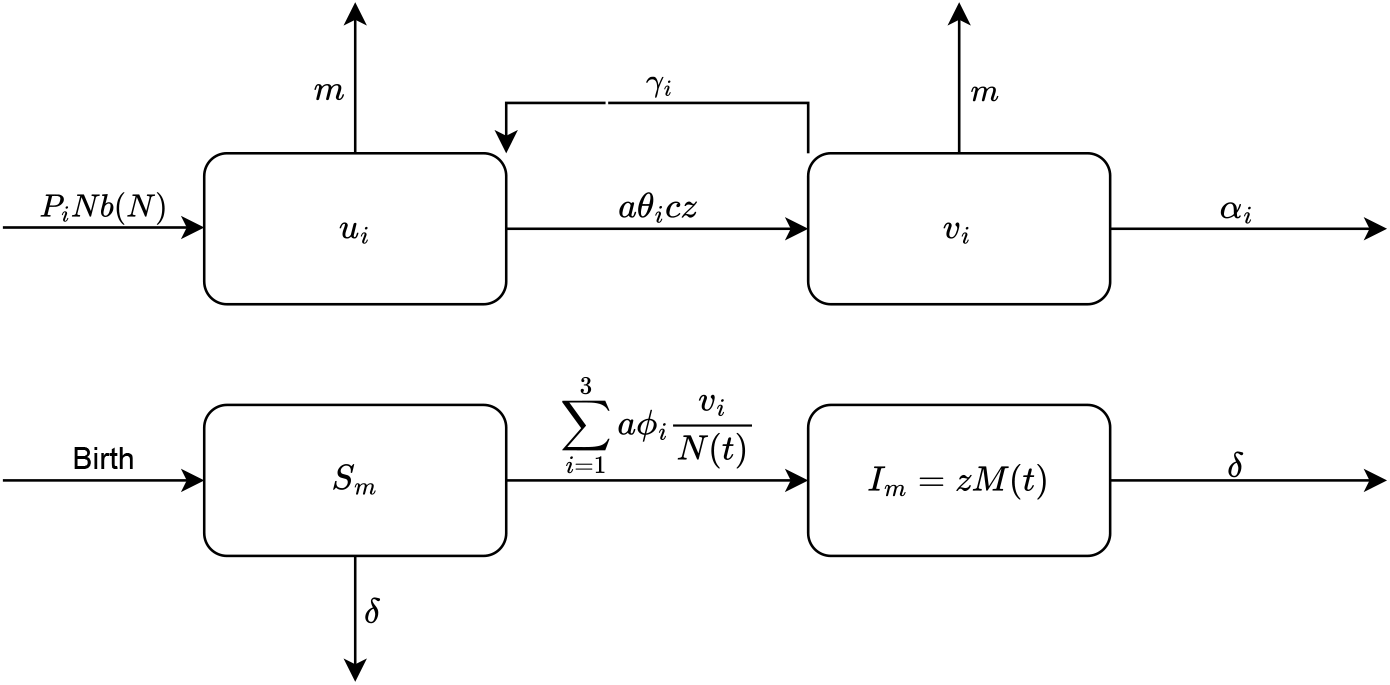
Flowchart of the population dynamics of humans (above) and mosquitoes (below), as described in (3).

The paper is structured as follows: we introduce the terms, parameters, and variables of our mathematical model. We then describe our model and its dynamics. We formulate the slow system, from which we deduct its manifold and the fitness of the the expressed Duffy allele. We use this information to simulate the fast system and to determine its next generation matrix, basic reproduction number, and equilibria. We conduct a global sensitivity analysis on the model, eventually including biting rate seasonality, and then estimate our parameters. We then conclude the paper by assessing the results and implications of our model.

## 2 Model

### 2.1 Model Formulation

There are four possible phenotypes of the Duffy antigen: Duffy-positive, where two of the alleles (FY*A and/or FY*B) alleles are present; the two heterozygous Duffy-positive traits, where one of the two alleles is present and the other is silent; and the homozygous Duffy-negative trait, where neither the FY*A or FY*B allele is expressed. For our paper, we use the following nomenclature for the Duffy blood group [11, 15, 17]:

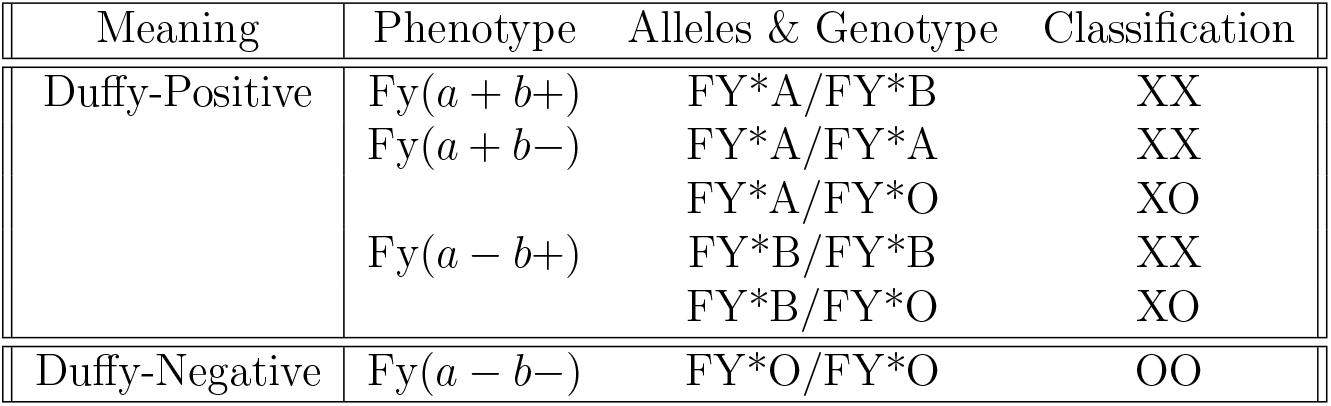

| Meaning | Phenotype | Alleles & Genotype | Classification |
| --- | --- | --- | --- |
| Duffy-Positive | Fy($a + b+$) | FY*A/FY*B | XX |
| | Fy($a + b-$) | FY*A/FY*A | XX |
|  |  | FY*A/FY*O | XO |
| | Fy($a - b+$) | FY*B/FY*B | XX |
|  |  | FY*B/FY*O | XO |
| Duffy-Negative | Fy($a - b-$) | FY*O/FY*O | OO |

In this paper, we use the term XX to refer to the presence of two expressed Duffy alleles, XO for the presence of one expressed Duffy allele and one silent Duffy allele, and OO for the presence of two silent Duffy alleles. The subscript *i* denotes the genotype: *i* = 1 for XX, *i* = 2 for XO, and *i* = 3 for OO.

We use Feng and Castillo-Chavez’s model [5] to describe the dynamics of the sickle cell trait. The difference between the models is that whereas the sickle cell model developed by Feng incorporates a death rate associated with the sickle cell trait, there is no death rate associated to the Duffy antigen trait. Furthermore, our model considers the impact of seasonality on our numerical simulations. The variables of our initial model are described in Table 1 and a summary of the parameters is presented in Table 2.1.

**Table 1.** Variables and description for original model. Our values for *P*_*i*_ and *w*_*i*_ are based in Rio Pardo, Brazil.

| State Variable | Description |
| --- | --- |
| $u_i$ | Uninfected individuals of genotype $i$ |
| $v_i$ | Infected individuals of genotype $i$ |
| $z$ | Fraction of infected mosquitoes |
| $x_i$ | Fraction of uninfected individuals of genotype $i$ |
| $y_i$ | Fraction of infected individuals of genotype $i$ |
| $w_i$ | Fraction of individuals in each genotype |
| $N$ | Total population |
| $S_m$ | Number of uninfected mosquitoes |
| $I_m$ | Number of infected mosquitoes |
| $M$ | Total number of mosquitoes |
| $P_1$ | Fraction of total births with XX genotype |
| $P_2$ | Fraction of total births with XO genotype |
| $P_3$ | Fraction of total births with OO genotype |
| $w_1$ | Fraction of XX people |
| $w_2$ | Fraction of XO people |
| $w_3$ | Fraction of OO people |
| $p$ | Frequency of the X-gene |
| $q$ | Frequency of the O-gene |

**Table 2.** Parameters, descriptions, values, units, and source..

| Human Parameters | Description | Value | Unit | Source |
| --- | --- | --- | --- | --- |
| $P_1$ | Fraction of total births with XX genotype | 0.6657 | people | [15] |
| $P_2$ | Fraction of total births with XO genotype | 0.3003 | people | [15] |
| $P_3$ | Fraction of total births with OO genotype | 0.0339 | people | [15] |
| $w_1$ | Fraction of XX people | 0.6623 | N/A | [15] |
| $w_2$ | Fraction of XO people | 0.3072 | N/A | [15] |
| $w_3$ | Fraction of OO people | 0.0304 | N/A | [15] |
| $p$ | Frequency of the X-gene | $w_1 + w_2/2$ | | |
| $q$ | Frequency of the O-gene | $1 - p$ | | |
| $N$ | Total Population | 690 | People | [15] |
| $b_0$ | Base human birth rate | 1.26e-5 | 1/day | [3] |
| $K$ | Density dependent reduction in birth rate | 10000 | people | [5] |
| $\theta_1$ | Probability of infection from infected bite for XX | 0.78 | N/A | [4] |
| $\theta_2$ | Probability of infection from infected bite for XO | 0.78 | N/A | [4] |
| $\theta_3$ | Probability of infection from infected bite for OO | 0.00702 | N/A | [4, 15] |
| $\gamma_1$ | Recovery Rate for XX | 1/24 | 1/day | [26] |
| $\gamma_2$ | Recovery Rate for XO | 1/24 | 1/day | [26] |
| $\gamma_3$ | Recovery Rate for OO | 1/24 | 1/day | [26] |
| $m$ | Natural human death rate | 3.78e-5 | 1/day | [3] |
| $\alpha_1$ | Disease-induced death rate for XX People | 0.0003 | Deaths/infection | [18] |
| $\alpha_2$ | Disease-induced death rate for XO People | 0.0003 | Deaths/infection | [18] |
| $\alpha_3$ | Disease-induced death rate for OO People | 0.0003 | Deaths/infection | [18] |
| Mosquito Parameters |  |  |  |  |
| $a$ | Biting Rate | 0.5 | 1/day | [23] |
| $c$ | Mosquitoes per Human | 2.3 - 4.36 | Mosquitoes/Human | [24] |
| $\phi_1$ | Probability a mosquito is infected from XX | 0.326 | N/A | [1] |
| $\phi_2$ | Probability a mosquito is infected from XO | 0.326 | N/A | [1] |
| $\phi_3$ | Probability a mosquito is infected from OO | 0.30 | N/A | [21] |
| $\delta$ | Mosquito Death Rate | 0.315 | 1/day | [2] |

Let *u*(*t*) be the total number of uninfected people and *v*(*t*) be the total number of infected people at time *t*. Thus, the total population is given by *N* (*t*) = *u*(*t*) + *v*(*t*). Furthermore, let the subscript *i* = 1 describe individuals with the Duffy genotype XX; *i* = 2 describe individuals with the XO genotype; and *i* = 3 describe individuals with the OO genotype. For example, *u*_1_ denotes the number of uninfected people whose Duffy genotype is XX, and *v*_2_ denotes the number of infected people with the XO genotype. Lastly, let *S*_*m*_(*t*) be the total number of susceptible mosquitoes and *I*_*m*_(*t*) be the total number of infected mosquitoes, where the total number of mosquitoes in the population is given by *M* (*t*) ≡ *S*_*m*_(*t*) + *I*_*m*_(*t*) at time *t*.

The mathematical model is derived based on the following assumptions:

1. New uninfected people of genotype *i* are born at the rate *P*_*i*_*Nb*(*N*), where *b*(*N*) represents the per capita birth rate as a function of the total population, and *P*_*i*_ is the fraction of births with genotype *i*. We define *b*(*N*) as 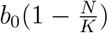, where *b*_0_ is the base birth rate and *K* is the carrying capacity. We assume that people die of non-disease-induced causes at the rate of *m*. Infection occurs in humans at the rate of *aθ*_*i*_*c*, where *c* is the mosquito-to-human ratio, *a* is the mosquito biting rate and *θ*_*i*_ is the probability a human becomes infected after being bitten by an infected mosquito. Lastly, infected individuals recover and enter the uninfected class at the rate *γ*_*i*_.
2. Individuals enter the infected compartment at a rate *aθ*_*i*_*c*. We assume individuals either die of natural causes at a rate *m*, or die due to malaria at a rate *α*_*i*_. Additionally, infected individuals may recover and leave the infected class at a rate *γ*_*i*_.
3. The mosquitoes are born into the susceptible class at rate *b*_*m*_. The term 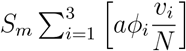 describes the transition from susceptible to infected mosquitoes, where *a* is the mosquito biting rate, and *ϕ*_*i*_ represents the probability that a mosquito becomes infected after biting an infected human of genotype *i*. Note that 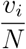 can be thought of as the probability that a mosquito bites a person of genotype *i* given that they bite an individual. Lastly, we assume both susceptible and infected mosquitoes die at the rate of *δ*. We then add the susceptible and infected mosquito classes to create the compartment 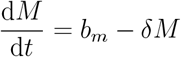, which we use to monitor the total population of mosquitoes.

Thus we have the system:

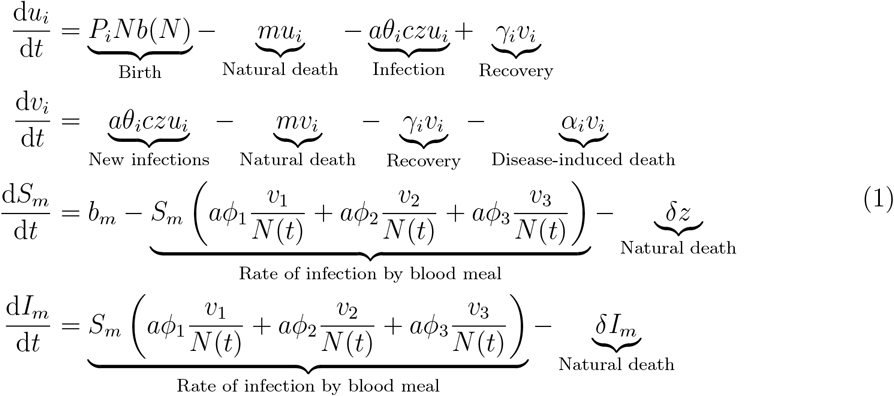

The flowchart below describes how people and mosquitoes move between their respective classes.

### 2.2 Human population genetic model

Our model requires distinguishing between the dynamics of each population’s genotype, because malaria dynamics occur on two time scales. Thus, we used Punnett squares to determine the fraction of total births of each genotype, in the Rio Pardo locality of Amazonas, Brazil, denoted by *P*_*i*_. We combine the expressed alleles FY*A and FY*B into a singular theoretical allele X. Likewise, we use O to represent the silent allele FY^ES^ (where ES means “erythrocyte silent”). Let *w*_*i*_ be the fraction of the total population by genotype (*w*_1_: frequency of XX, *w*_2_: frequency of XO and *w*_3_: frequency of OO).

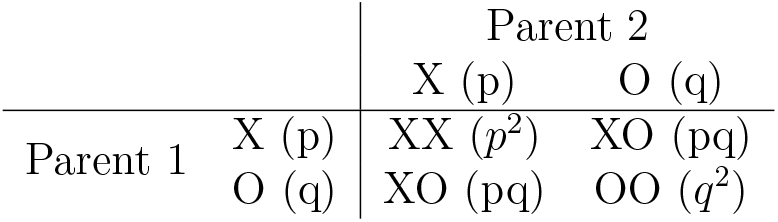

The previous table shows how we track the gene frequencies of X and O. Thus, *p*^2^+2*pq*+*q*^2^ = 1 so let’s redefine *P*_*i*_ as follows (note *w*_3_ = 1 − *w*_1_ − *w*_2_):

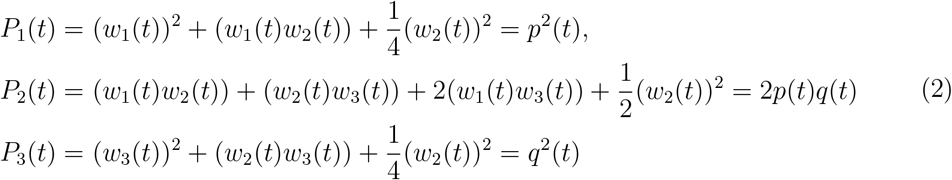

The assumptions above lead to the following nonlinear ordinary differential equations, which model the disease dynamics of each genotype:

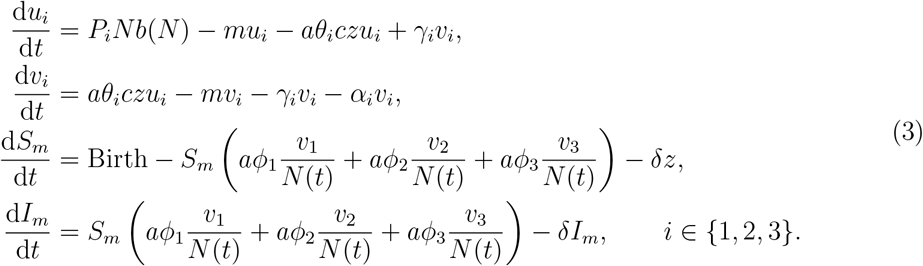

### 2.3 Re-scaled model

The re-scaling of the model is done for the sake of simplification. By re-scaling our original model, we can use the proportion of genotypic populations in Rio Pardo, Brazil, and apply the model to different populations in other regions. After this, perturbation analysis allows us to separate the fast and slow dynamics in our system. As infection happens on a much faster scale than human birth, death and evolution, it is crucial to create this separation so we can determine the fitness of our genotypes, and to study the rate of infection by genotype. Let 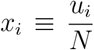 and 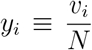 to represent the fraction of people with genotype *i* who are uninfected and infected, respectively. Thus, *w*_*i*_ ≡ *x*_*i*_ + *y*_*i*_ is the fraction of the population with genotype *i*. We define 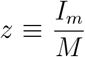 as the fraction of mosquitoes that are infected. Lastly, we let 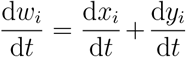 and substitute 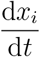 and 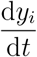, similarly, 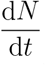 is found using the relation 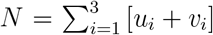. For further simplification, we define *β*_*hi*_ = *aθ*_*i*_*c* to replace the infection rate of humans after being bitten by mosquitoes, and *β*_*vi*_ = *aϕ*_*i*_ to replace the infection rate of mosquitoes after biting humans.We then have the following re-scaled model:

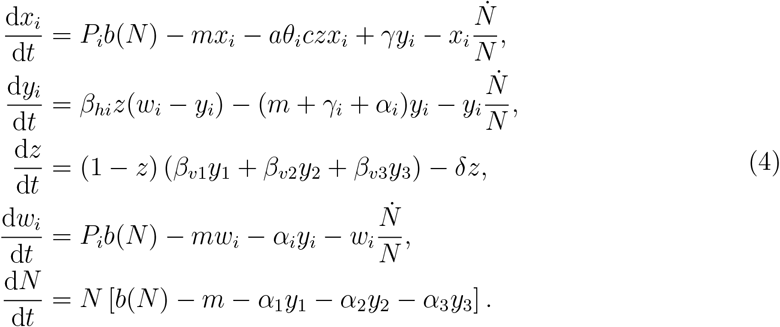

### 2.4 Fast Dynamical System

The *fast system*, which describes short-term disease dynamics, uses *t* as the time variable applied to disease transmission. Whereas the slow system operates in a matter of decades depending on the value of *ϵ*, the fast system operates on a more limited scale in a matter of days. This neglects birth rates and death rates, and instead focuses on *P. vivax* transmission and recovery. Let *ϵ >* 0 be small, and let 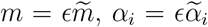 and 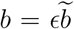. Substituting into system (4) yields:

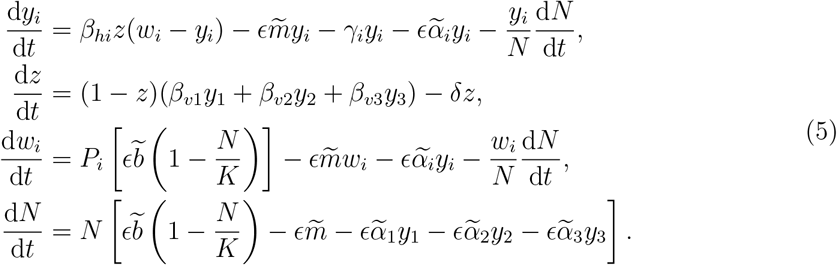

In the fast system, the parameter *ϵ* approaches zero so that the birth and death rate are negated. Because 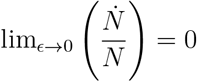, then as *ϵ* → 0, the fast system then becomes

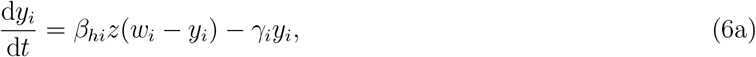

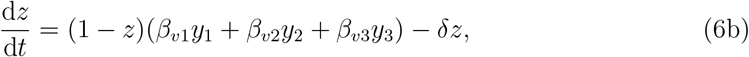

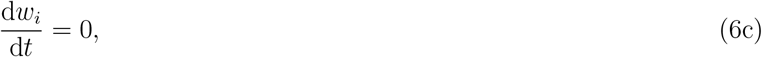

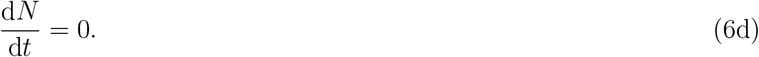

### 2.5 The Next Generation Matrix and the Basic Reproduction Number of the Fast System

The next generation matrix is used to calculate the *basic reproduction number*, which predicts the number of *P. vivax* cases caused by one preceding incidence. We use the remaining equations in the fast system to form the initial matrices

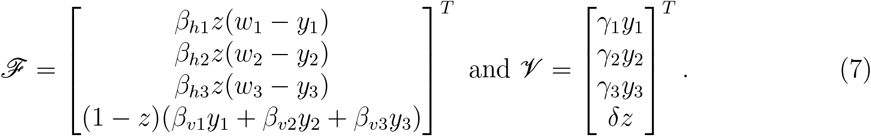

Taking the Jacobian of ℱ and *V*, respectively, yields:

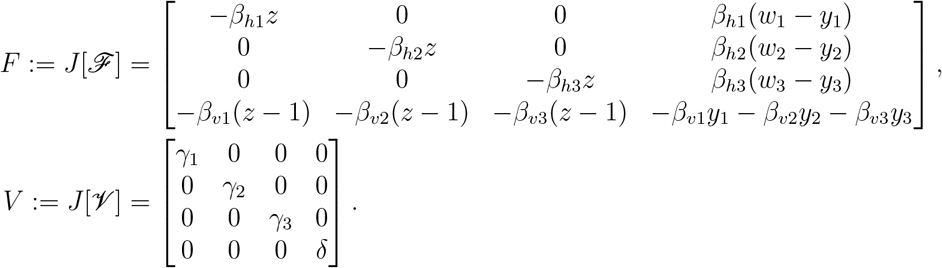

Thus, at the disease free equilibrium *E*_0_ where (*y*_1_, *y*_2_, *y*_3_, *z*) = (0, 0, 0, 0), the next generation matrix is:

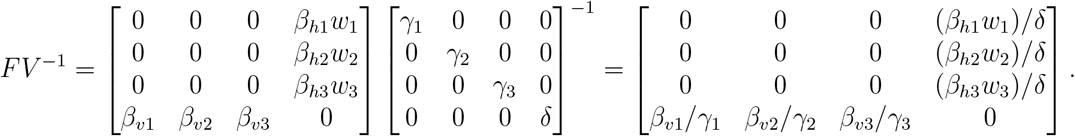

The spectral radius of the next generation matrix is our basic reproduction number, denoted by ℛ_0_.

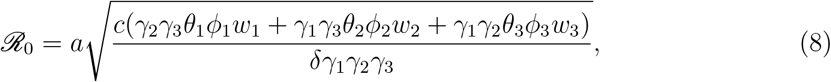

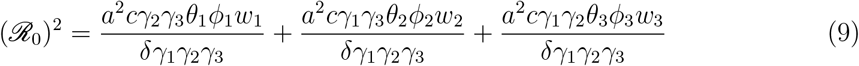

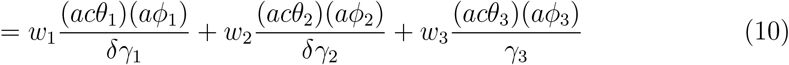

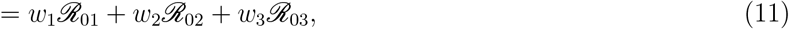

where 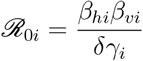. For convenience, we define

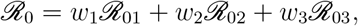

which doesn’t affect the validity of the inequalities ℛ_0_ *>* 1 and ℛ_0_ *<* 1.

### 2.6 Conditions for Stability of Disease-Free Equilibrium

**Theorem 1** *The disease free equilibrium E*_0_ *is locally asymptotically stable if* ℛ_0_ *<* 1 *and unstable if* ℛ_0_ *>* 1.

*Proof*. To determine the stability of *E*_0_, we first find the Jacobian of the *fast system* and evaluate it at *E*_0_.

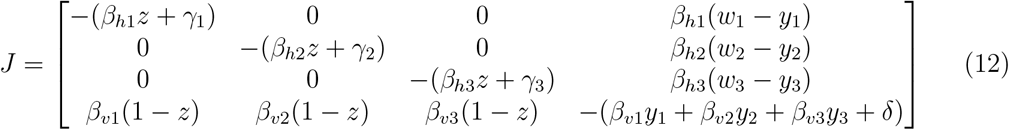

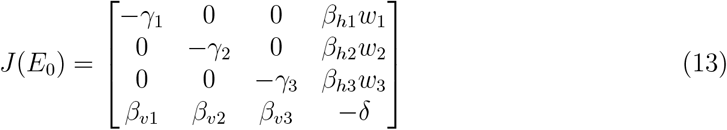

Note that *J*(*E*_0_) can be expressed as *M* − *D* where *M* and *D* are both non-negative matrices and *D* is additionally a diagonal matrix.

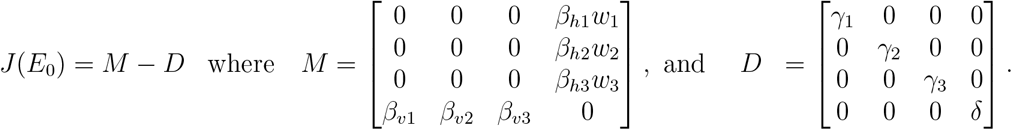

From here, we use the following theorem.

**Theorem 2** *Let M and D be matrices such that every entry is nonnegative and D is a diagonal matrix and let H* = *M* − *D. If the dominant eigenvalues of MD*^−1^ *are less than* 1, *then the real part of the eigenvalues of H are less than* 0. *[5]*

It can be shown that the matrix *MD*^−1^ has two eigenvalues of 0 and the remaining two are

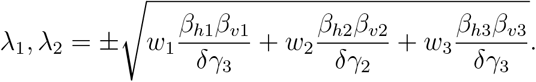

Making the substitution 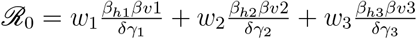 yields

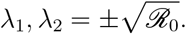

Note that 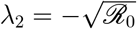 is always less than 1 since ℛ_0_ is always greater than or equal to 0. Additionally, *λ*_1_ *<* 1 when ℛ_0_ *<* 1 and *λ*_1_ *>* 1 when ℛ_0_ *>* 1. By Theorem 2, the real part of all the eigenvalues of *J*(*E*_0_) are negative when ℛ *<* 1, meaning the disease-free equilibrium is locally asymptotically stable when ℛ_0_ *<* 1. Likewise, *E*_0_ is unstable when ℛ_0_ *>* 1. □

### 2.7 Location of Nontrivial Equilibrium

The nontrivial equilibrium 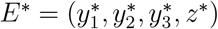 is a solution to the following system.

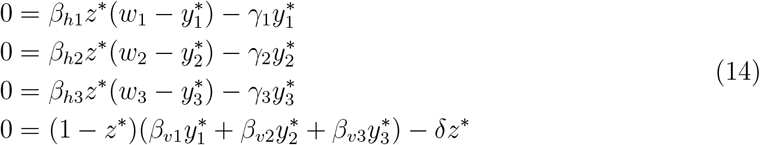

For convenience, we define 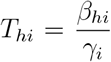. Substituting and simplifying yields the following expression for 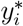:

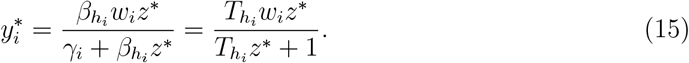

To find the solutions for *z*^∗^, we substitute out 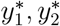, and 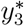 in (14) and divide out the trivial solution of *z*^∗^ = 0. Additionally, we let *w* = *w*_1_ + *w*_2_ and assume *T*_*h*1_ = *T*_*h*2_ and ℛ_01_ = ℛ_02_. This yields the quadratic:

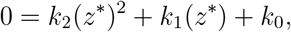

where

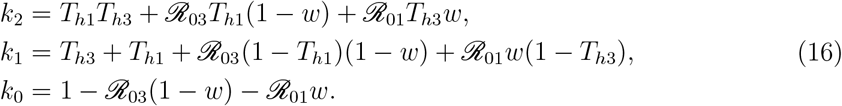

Note that *k*_0_ can be rewritten as *k*_3_ = 1 − ℛ_0_, and let *h*(*z*) = *k*_2_(*z*)^2^ + *k*_1_(*z*) + *k*_0_. Note that *z*^∗^ is by definition a solution to the equation *h*(*z*) = 0.

#### Lemma 1

*If* 0 *<* ℛ_0_ *<* 1 *then k*_1_ *>* 0.

*Proof*. Let 0 *<* ℛ_0_ *<* 1 and define *T*_0_ = max(*T*_*h*1_, *T*_*h*3_). Based on these two statements,

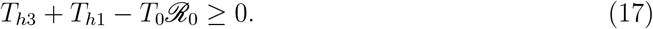

Additionally, since *T*_0_ ≥ *T*_*h*1_ and *T*_0_ ≥ *T*_*h*3_, we know

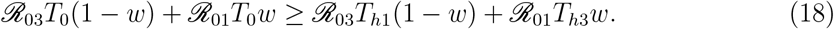

From our definition of *k*_1_ (16),

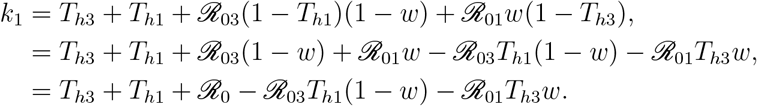

From here, using relation (18) yields

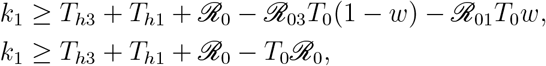

and relation (17) implies

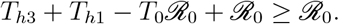

Combining the two results above yields *k*_1_ ≥ ℛ_0_. Therefore, if 0 *<* ℛ_0_ *<* 1, then *k*_1_ *>* 0.

**Theorem 3** *Assuming all parameters are biologically possible, the fixed point E*_0_ *in the* ***fast system*** *(6a) is biologically impossible when* ℛ_0_ *<* 1 *and biologically possible when* ℛ_0_ *>* 1.

*Proof of Theorem 3*. Let all parameters be biologically possible and 0 *<* ℛ_0_ *<* 1. From Equation 16 it follows that *k*_2_ *>* 0 and *k*_0_ *>* 0, due to the respective conditions above. Additionally, Lemma 1 implies, *k*_1_ *>* 0.

Let 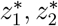 be the two solutions to the quadratic *h*(*z*) = 0. Then, by Vieta’s formulas, the solutions to *h*(*z*) = 0 satisfy

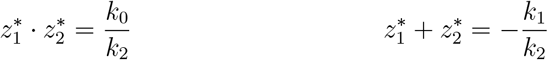

and using the conditions *k*_0_, *k*_1_, *k*_2_ *>* 0, they simplify to

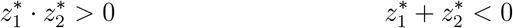

The inequality on the left rules out the possibility that exactly one of the roots of *h*(*z*) is positive and the inequality on the left implies that the roots cannot both be positive. Therefore, neither 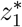 or 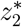 are positive. As such, when 0 *<* ℛ_0_ *<* 1, the nontrivial equilibrium is not biologically possible since *E*^∗^ would require a negative or complex fraction of mosquitoes to be infected, neither of which are possible in the real world.

To prove the second part of Theorem 3, let all parameters be biologically possible and let ℛ_0_ *>* 1. Similar to earlier, these two conditions imply *k*_2_ *>* 0, *k*_1_ *<* 0 and *k*_0_ *<* 0. Similar to before, applying these conditions to Vieta’s Formulas yields

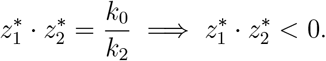

Since *k*_1_, *k*_2_, *k*_3_ are all real numbers, 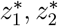 must also, so the inequality above must imply that either 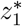 or 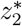 is positive, but not both. Let *z*^∗^ be this unique positive solution to *h*(*z*) = 0.

Next, we wish to show that 0 *< z*^∗^ *<* 1. Since ℛ_0_ *>* 1, we know *k*_0_ *<* 0 by Lemma 1. Additionally,

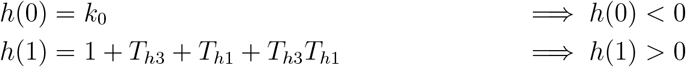

since *T*_*h*1_ and *T*_*h*3_ are greater than 0 when all parameters are biologically possible. Because *h*(*z*) is a continuous function with *h*(0) *<* 0 and *h*(1) *>* 0, the Intermediate Value Theorem implies that there exists a solution to *h*(*z*) = 0 between *z* = 0 and *z* = 1. Since *z*^∗^ is the only positive solution to *h*(*z*) = 0, we conclude that

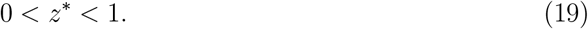

Recall that *z*^∗^ represents the fraction of mosquitoes at the equilibrium point *E*^∗^. As such, *z*^∗^ is biologically possible when ℛ_0_ *>* 1 since 0 *< z*^∗^ *<* 1.

Lastly, we wish to show that 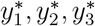 are also between 0 and 1 when all parameters are biologically possible. Recall that Equation 15 tells shows

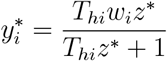

Since *w*_*i*_ represents the fraction of the population with genotype *i, w*_*i*_ must satisfy 0 ≤ *w*_*i*_ ≤ 1 in the real world. As such, the denominator for 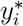 is necessarily greater than the numerator. Therefore, at the nontrivial equilibrium, 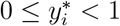, meaning 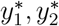, and 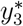 are all biologically possible. As such, we conclude that *E*^∗^ is biologically possible when all parameters are biologically possible and ℛ_0_ is greater than 1.

### 2.8 Conditions for Stability of Endemic Equilibrium

In the previous section, we proved that the endemic equilibrium *E*^∗^ exists in biologically possible space only when ℛ_0_ *>* 1. We now present the following theorem on the stability of *E*^∗^.

**Theorem 4***If* ℛ_0_ *>* 1 *and all parameters are biologically possible (i*.*e. E*^∗^ *is biologically possible), then the fixed point E*^∗^ *is stable*.

*Proof*. Taking the Jacobian of the *fast system* (12) and evaluating it at *E*^∗^ yields

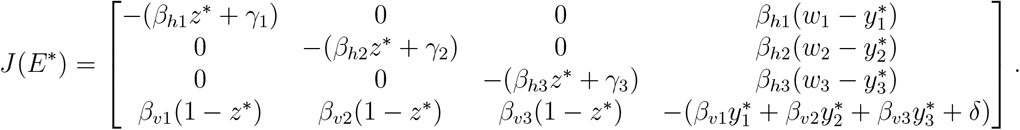

Note that *J*(*E*^∗^) = *M* − *D* where *M* and *D* are both non-negative matrices and *D* is a diagonal matrix.

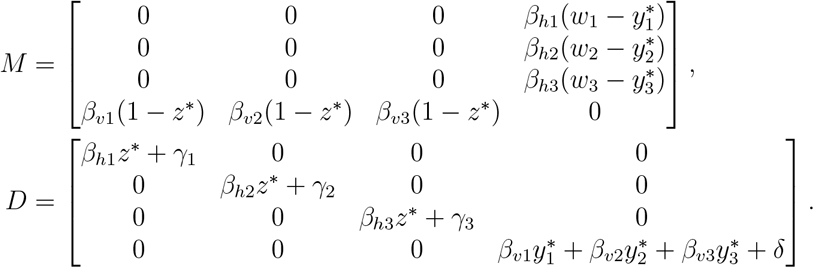

Recall that Theorem 2 states that the real part of the eigenvalues of *J*(*E*^∗^) are negative if the dominant eigenvalue of *MD*^−1^ is less than 1.

It can be shown that *MD*^−1^ has a double eigenvalue of 0 and the other two are

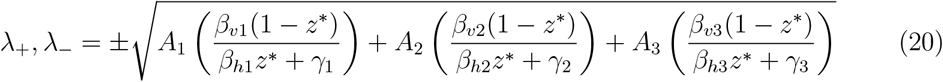

where

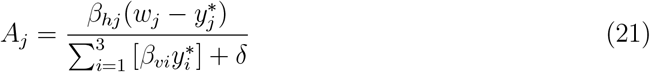

Note that *λ*_−_ is always less than 1 provided that all parameters are within biologically possible bounds. Next, we wish to show that *λ*_+_ is also less than 1.

From Equation 14, recall that

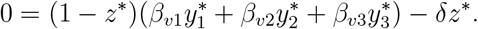

Manipulating this equation to isolate *z*^∗^ yields

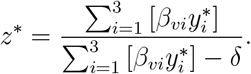

Using the equations of form 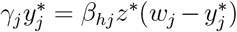 from Equation 14, we obtain the following expression for *z*^∗^.

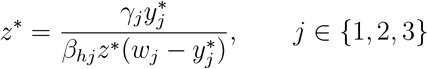

Setting these two expressions for *z*^∗^ equal and multiplying both sides by the fraction 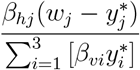 yields

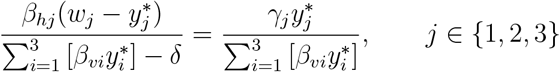

Note that the left side of this expression is equivalent to our definition of *A*_*j*_, Equation(21). As such,

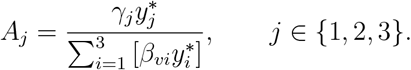

Furthermore,

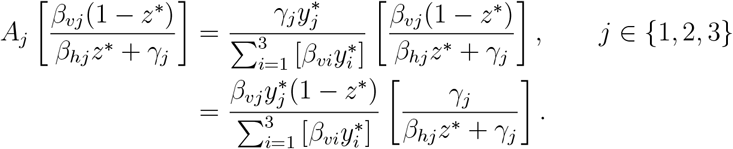

From here, since all parameters are assumed to be biologically possible, *β*_*hj*_ + *γ*_*j*_ *> γ*_*j*_. Using this inequality, we find

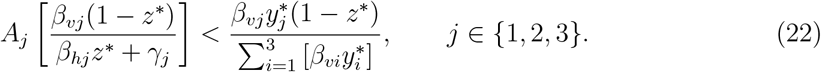

Next, we sum Equation (22) over *j* = 1, 2, 3 to obtain the following. Note that the left side is equivalent to (*λ*_+_)^2^ by (20).

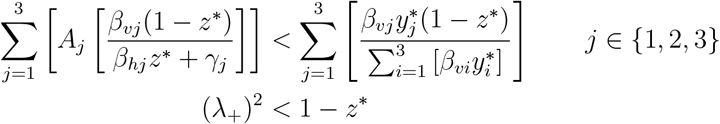

Recall from Equation (19) that 0 *< Z*^∗^ *<* 1. Therefore *λ*_+_ *<* 1. Since both *λ*_−_ and *λ*_+_ are less than 1, Theorem 2 applies, meaning that the real part of the eigenvalues of *J*(*E*^∗^) are negative. As such, we conclude that *E*^∗^ is stable if ℛ_0_ is greater than 1 and all parameters are biologically possible. □

### 2.9 Slow Dynamical System

We use a fast-slow decomposition to analyze the dynamics of our model. The *slow system* employs the time-scale *τ* = *ϵt*, for some small positive value *ϵ*. As such, large changes in *t* result in small changes to *τ*, making analysis of the slow system well suited to investigate dynamics which unfold over long periods of time, such as evolution. Model (4) provides a template for modeling the slow and fast dynamics of malaria transmission in the Rio Pardo. In the slow system, time is parameterized to be measured in *τ* = *ϵt*. From this equation, we determine that

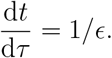

By letting 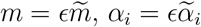 and 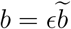 we obtain the following,

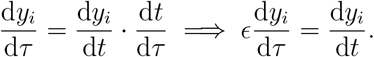

A similar pattern applies to all derivatives with respect to *t*. In this fashion, the original model of the system is simplified to the slow system as follows:

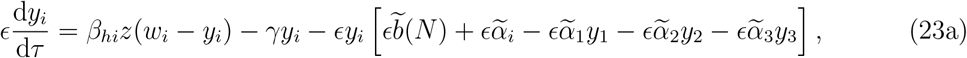

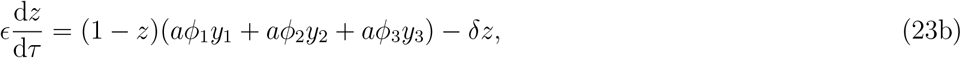

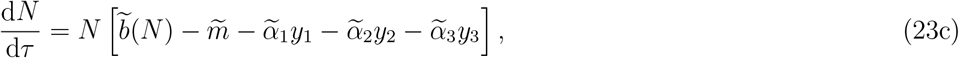

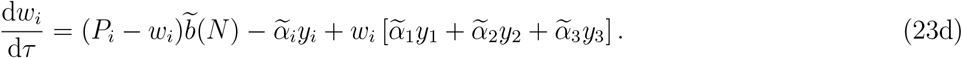

#### 2.9.1 Manifold of Slow System

This system has a four-dimensional slow manifold:

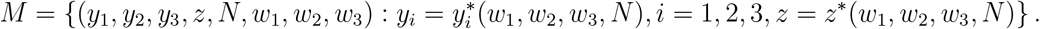

The functions 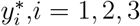 and *z*^∗^ are given in (15) and (16). The slow dynamics on *M* are described by the following equations:

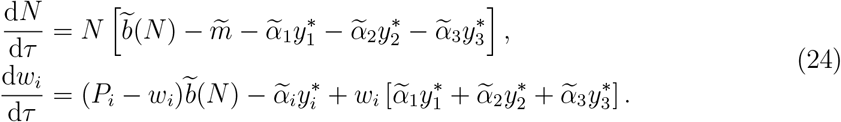

#### 2.9.2 Determining Fitness of the X-gene

The fitness of the X gene, that being the presence of the *Fy*^*a*^ and/or *Fy*^*b*^ antigens in reticulocytes, determines the capacity of the Duffy-positive gene to be passed down to future generations. By calculating the fitness of the X gene, we determine the influence of selection on the prevalence of the Duffy-negative trait in our region of study. We denote the frequency of the X-gene by 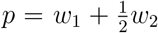, where *w*_1_ is the frequency of XX individuals and *w*_2_ is the frequency of XO individuals. The invasion ability of the X-gene is described by

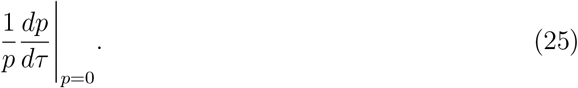

We will use this quantity to define the fitness of the X-gene, ℱ. We assume all genotypes have the same death rate, *m*. We define the following out of convenience:

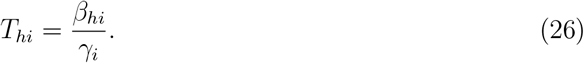

Using equations from the slow system dynamics for *w*_*i*_ we obtain

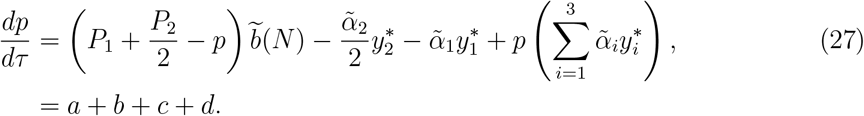

We have that 0 ≤ *w*_1_, *w*_2_ ≤ *p*. So, when *p* = 0, we must have that *w*_1_ = *w*_2_ = 0 and subsequently *w*_3_ = 1. Substituting the expressions for *P*_1_ and *P*_2_ in (2) in (27) we obtain:

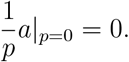

Assume 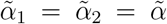 and 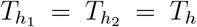. Note also that 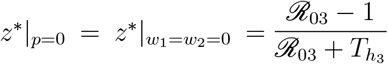. Then,

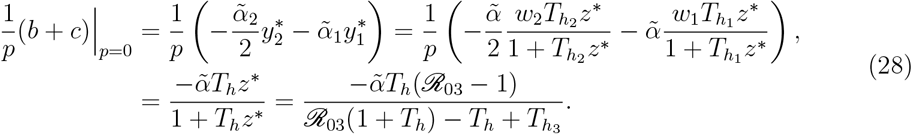

Further, given that *y*_1_|_*p*=0_ = 0, *y*_2_|_*p*=0_ = 0 and

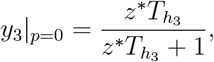

we have that

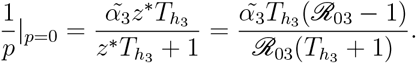

Combining the terms yields

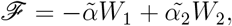

where 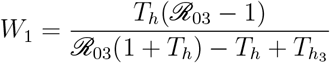 and 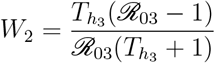.

We plot the fitness of the *X*-gene utilizing the parameters obtained from the literature in Fig. 4. Notice that as the infection rate for *OO* people decreases, the fitness or the invasion ability of the *X* gene decreases. Also, notice that as the recovery rate for *OO* people increases, the fitness of the *X* gene decreases meaning the *X* gene invades less.

**Figure 4.**
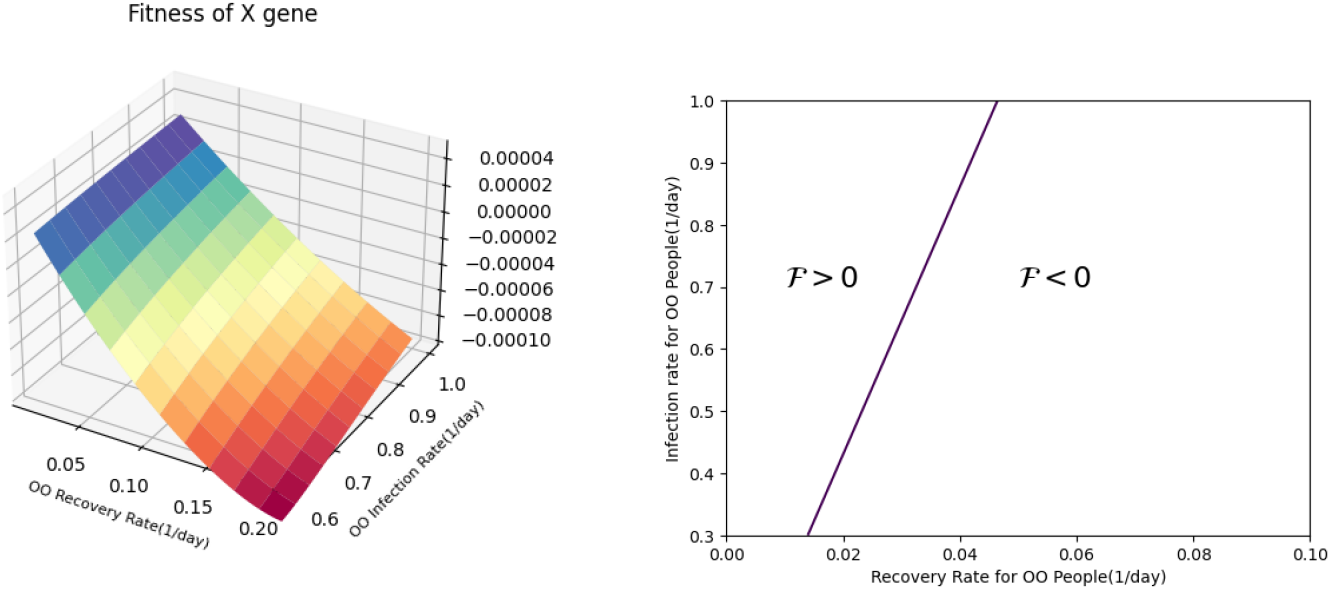
The left plot shows the fitness of the X gene plotted with respect to *γ*_3_ (recovery rate for Duffy-negative people) and *β*_*h*3_ (infection rate for Duffy-negative people).The right plot is the contour plot that shows the boundary where the fitness is greater or less than zero or rather when the X-gene invades.

## 3 Sensitivity Analysis

We conduct global sensitivity analysis on our model to determine which parameters are most influential to the model’s behavior. We include biting rate seasonality on our global sensitivity analysis of the fast subsystem, as outlined by the inter-compartmental approach from Renardy et al [20].

There are multiple methods to perform global sensitivity analysis. For nonlinear and monotonic relationships, measures that work the best are Spearman rank correlation coefficient (RCC or Spearman’s *ρ*), partial rank correlation coefficient (PRCC), and standardized rank regression coefficients (SRRC) [16]. The Sobol method is effective for nonlinear non-monotonic trends, as do the Fourier amplitude sensitivity test (FAST) and its extended version (eFAST). The derivative based one at a time (OAT) method can be used on any continuous system where it isn’t computationally difficult to calculate the partial derivatives. Since the relationship in the fast subsystem with seasonality is nonlinear and non-monotonic with computationally possible derivatives we can either utilize the eFAST or the OAT method. We proceed with the OAT method to determine the elasticity of ℛ_0_ and conduct eFAST to determine the sensitivities of parameters of the full system.

### 3.1 Calculating The Elasticity of *R*_0_

We compute the sensitivity indices of the parameters to determine how much each parameter influences ℛ_0_. The sensitivity index for a parameter *p* is calculated as

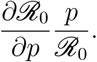

The sensitivity indices are as shown in Figure 5. Since the biting rate *a* and the mosquito death rate *δ* rely on the population of mosquitoes, ℛ_0_ is most drastically influenced by these parameters. The parameters *θ*_3_, *ϕ*_3_, and *γ*_3_ all have low elasticities because of the low percentage of the population with absence of the Duffy trait.

**Figure 5.**
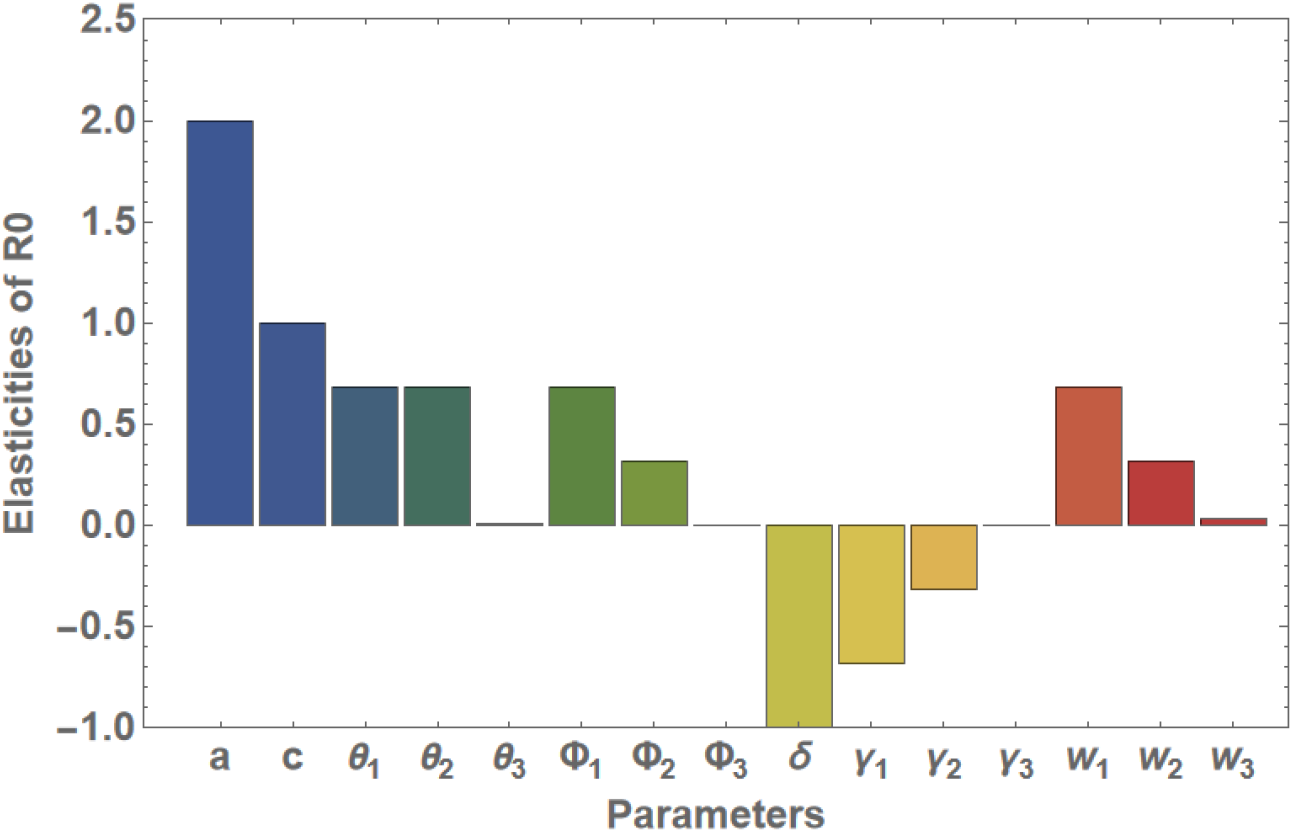
Sensitivity indices for *R*_0_ for each parameter in Rio Pardo

### 3.2 Global Sensitivity Analysis on the Full System

We compute the first order and total order sensitivity indices of each parameter in the Full System(Equation 4) with respect to the total fraction of people infected, *y* which is *y*_1_ + *y*_2_ + *y*_3_ utilizing eFAST. The eFAST sensitivity method measures the fraction of output variance that are affected by variance in the input parameters. Then, statistical inference is run via the use of a dummy parameter that decides which value of the sensitivity index is significantly different from zero.[20] The initial parameter values used were obtained after conducting parameter estimation, fitting the full system to time series data of cases in Brazil [8]. See the table 3 from the Parameter Estimation Section. The upper and lower bounds of the parameters were found by taking 80% and 120% of the initial estimated values. 280 samples were used to generate the sensitivity indices. The results can be seen in Figure 6. An asterisk(*) above the bars means the variable has a p-value ≤ 0.05 indicating the variable is statistically significant. The parameter *a* associated with the mosquito bite rate has the highest sensitivity indices which lines up with the elasticities of ℛ_0_.

**Table 3.**
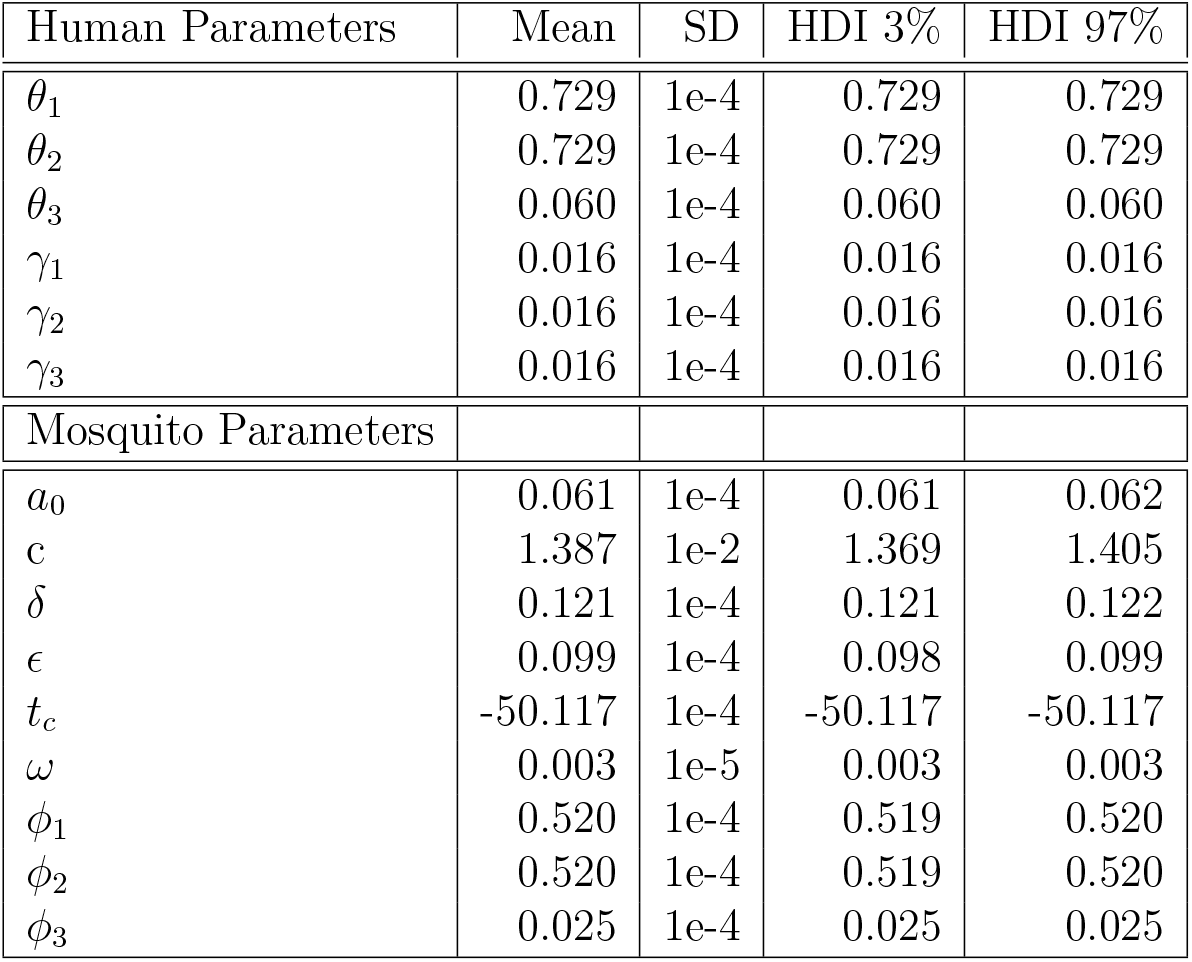
Estimated Parameters for Fast System with Biting Rate Seasonality.

**Figure 6.**
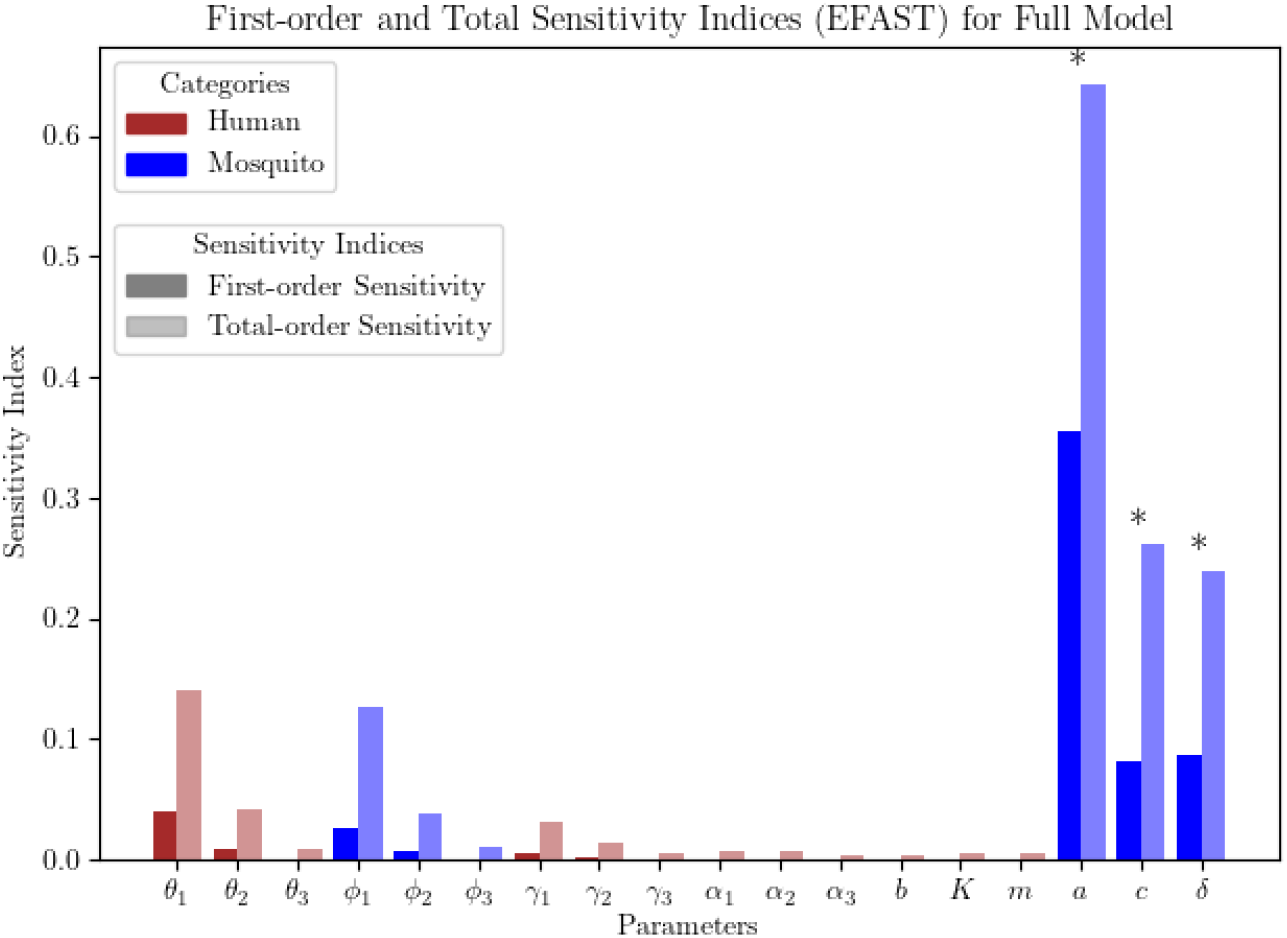
Total and First Order Sensitivity indices resulting from eFAST on Full Rescaled System. Significance (*p* ≤ 0.05) is indicated with an asterisk(∗)

## 4 Parameter Estimation

To estimate the transmission rates of malaria between mosquitoes and humans, we conduct parameter estimation on the fast subsystem using incidence data from Brazil and by assuming that Rio Pardo’s Duffy genotype frequency is representative of the wider Brazilian Amazon.

We begin by splitting up time series of cases in Brazil from August 2020 to May 2023 from Garcia et al. [8] to the number of people in each DARC genotype infected.

### 4.1 Relative Risk of Infection Based on Genotype

Due to the limitations of available literature, we use our known information to infer the infection rates of *P. vivax* malaria, based on the Duffy polymorphisms in Rio Pardo. For our model, we attempt to weight the relative risk of being infected with *P. vivax* malaria (*n*_*i*_) by an individual’s genotype. We presume that if everyone in the population contracts malaria at the same rate, then 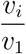 is equal to 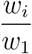. We used the values of variables provided in the paper by Kano et al. [15] to adjust the risk according to the genotype of each population, resulting in this system of equations:

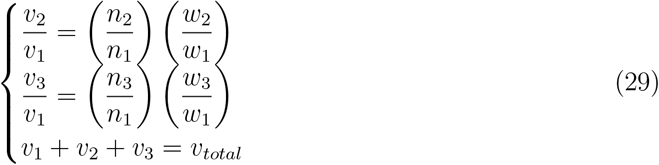

where *n*_*i*_ is the relative risk of infection of *P. vivax* malaria for someone with a genotype *i*. By calculating 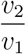 and 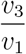 and solving the system of equations, we can determine the proportion of the infected population by genotype.

The values of *n*_*i*_ must be calculated to determine the relative risk of infection for each genotype. As determined by Kano et al. [15], let *n*_*aa*_ be the risk of infection due to the FY*A/FY*A genotype; *n*_*bb*_ be the risk of infection due to the FY*B/FY*B genotype:

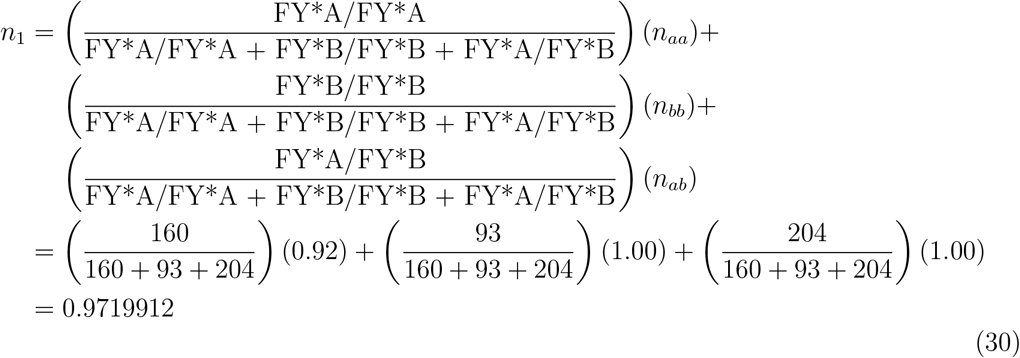

We define *n*_2_ as the weighted average of the relative risk of infection a heterozygous genotype with a silent allele. Let *n*_*ao*_ be the risk of infection for FY*A/FY*O genotype, and *n*_*bo*_ be the risk of the FY*B/FY*O genotype.

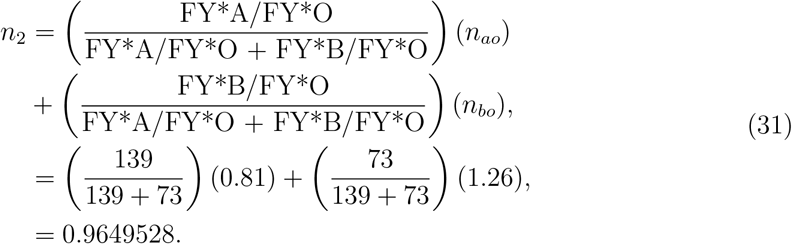

The relative risk 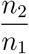 of the heterozygous genotype is therefore calculated as follows:

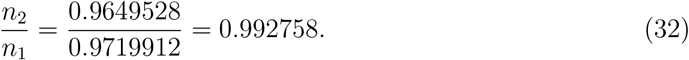

Thus, from Equation 29 we solve for 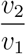 and 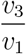:

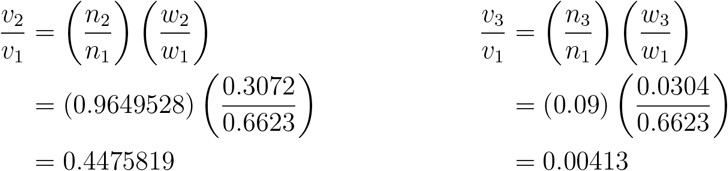

where 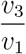 represents the homozygous Duffy-negative genotype.

### 4.2 Fitting the Fast System with Seasonality

The elasticity of ℛ_0_ and sensitivty of *a* in the full system substantiates that the mosquito biting rate is the most influential parameter to ℛ_0_. We can then account for the seasonality of biting rate by equating *a* to a sinusoidal model.

We begin the parameter estimation by incorporating the sinusoidal seasonality of the mosquito biting rate by including the below model from [14] to the biting rate of mosquitoes:

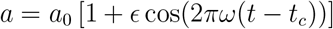

where *a*_0_ is the average biting rate, *ϵ* is the degree of variation around the biting rate, *ω* is the frequency of seasonal malaria epidemics and *t*_*c*_ is the phase shift in periodicity.

The Sequential Monte Carlo(SMC) method was used to conduct parameter estimation and determine the confidence intervals for parameters fitted. Specifically, the following quantity was minimized:

(Incidence of Total Cases(Data) - Incidence of Total Cases(Model)) +

(Incidence of XX Cases(Data) - Incidence of X Cases(Model)) +

(Incidence of XO Cases(Data) - Incidence of XO Cases(Model))

(Incidence of OO Cases(Data) - Incidence of OO Cases(Model)).

In Fig. 7, the points representing the data look very similar because they differ by a constant factor due to our assumptions when disaggregating the incidence data. We assumed that the distribution of people with each DARC genotype was equal to the distribution in Rio Pardo. Each parameter and initial condition value took on a Truncated Normal Distribution except sigma the standard deviation of the observations takes on a Half Normal Distribution. The results of the parameter estimation can be seen in Table 3 and Table 4 where SD refers to the standard deviation of each individual distribution. HDI 3% refers to the 3% high density interval or confidence interval and similarly for HDI 97%.

**Table 4.** Estimated Initial Conditions for Fast System with Biting Rate Seasonality.

| Initial Conditions | Mean | SD | HDI 3% | HDI 97% |
| --- | --- | --- | --- | --- |
| $u_1$ | 15840000.010 | 0.1 | 15839999.930 | 15840000.094 |
| $u_2$ | 7199999.996 | 0.1 | 7199999.914 | 7200000.074 |
| $u_3$ | 719999.964 | 0.1 | 719999.901 | 720000.032 |
| $v_1$ | 121.519 | 0.1 | 121.329 | 121.704 |
| $v_2$ | 60.586 | 0.1 | 60.411 | 60.775 |
| $v_3$ | 9.017 | 0.1 | 8.871 | 9.165 |
| $z$ | 0.002 | 1e-4 | 0.002 | 0.002 |

**Figure 7.**
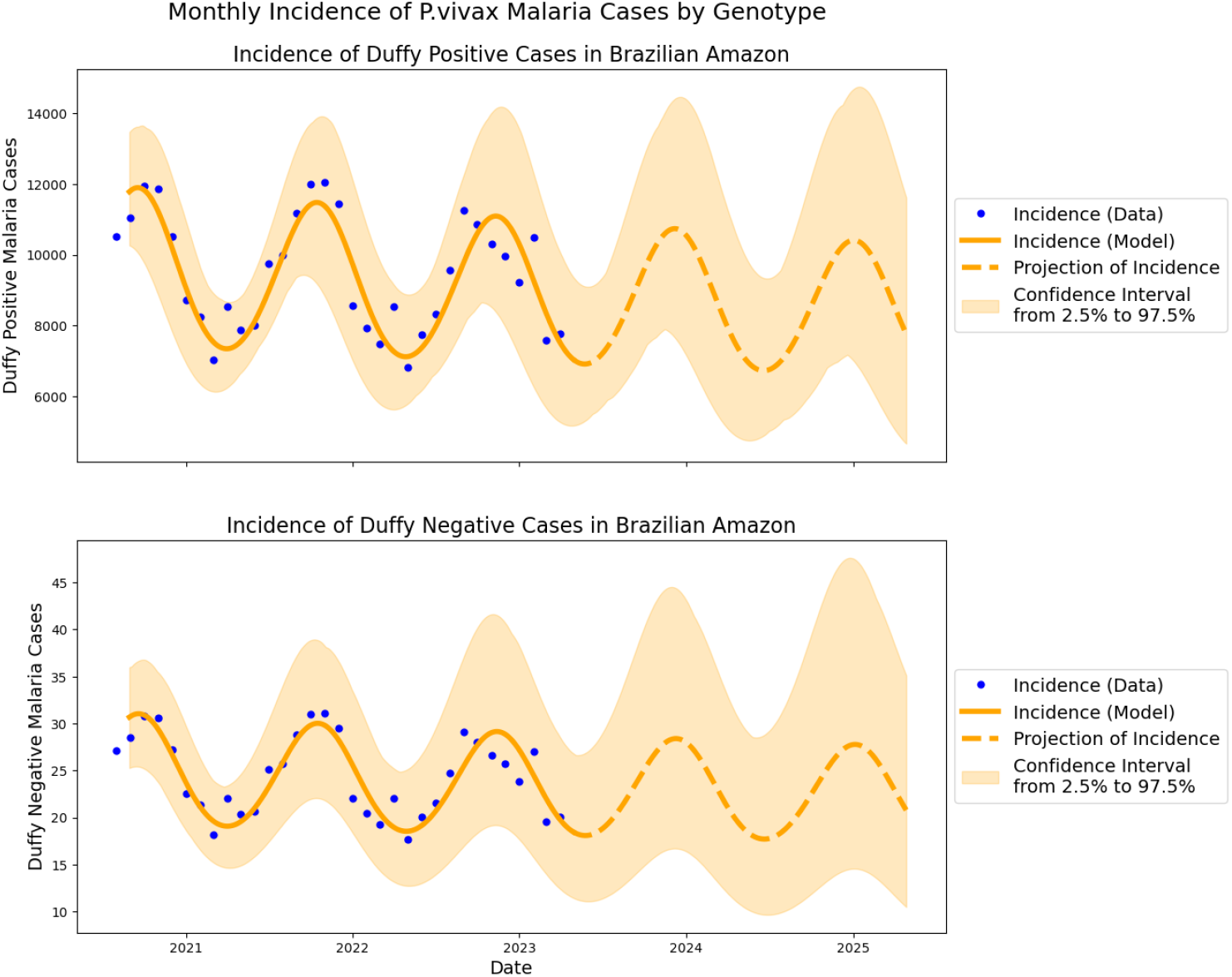
Fit of model to the estimated breakdown of *P. vivax* by Duffy positivity in the Brazilian Amazon

## 5 Results

We developed a multi-scale mathematical model that quantifies the impact of the DARC genotype on the epidemiology of *vivax* malaria, and the impact of *vivax* malaria on the long-term frequency of Duffy-positivity. From our sensitivity analysis of ℛ_0_ for the fast system without seasonal biting rate, we find that the mosquito biting rate, *a*, has the highest elasticity index, and therefore the largest impact on the value of ℛ_0_. Thus, varying the mosquito biting rate heavily influences the dynamics of *P. vivax*.

Since malaria incidence data exhibits seasonal fluctuations, we implemented seasonality into our model by making the biting rate a sinusoidal function of time. Using this updated model, we then conducted sensitivity analysis on the model’s prediction of the total number of people infected across genotypes. However, because different places have different percentages of the population with each genotype, we conducted this analysis using the genotype fractions from Macapa and Rio Pardo, where about 59% and 3% of people are Duffy-negative, respectively.

When comparing the results of the sensitivity analysis on the model incorporating seasonality, we find that in both Macapa and Rio Pardo, the recovery rates *γ*_1_ and *γ*_2_ have the largest impact on the number of total infected people, which reflects our model’s assumption that people leave the infected class at these rates. Aside from these two parameters, the average biting rate *a*_0_ has the largest sensitivity index, with *c*, the ratio of mosquitoes to human, coming in second. The population of mature female mosquitoes per area is thereby a driving force of infection by *P. vivax*. Given that the life cycle of mosquitoes is dependent on temperature and humidity, we defined the biting rate as a sinusoidal function. By fitting this to our data, we assume that biting rate is the diving force of this cycle. Our sensitivity analysis confirms this positive correlation between the biological reality of the vector and our model.

Similarly, mosquitoes are infected by taking a blood meal from human populations; however, taking this blood meal from a Duffy-negative individual has no effect on the rate of mosquito infection. Indeed, only *ϕ*_1_ and *ϕ*_2_ have positive sensitivity indices in both areas. In Rio Pardo, *w*_3_ has a negligible sensitivity index due to the small percentage of Duffy-negative people in Rio Pardo. In Macapa, *w*_3_ has greater sensitivity due to the larger percentage of Duffy-negative people in Macapa. However, *w*_3_ is still smaller than *w*_1_ and *w*_2_ due to the reduced transmission rates for Duffy-negative people.

We conducted parameter estimation on the fast system by utilizing a time series of cases in Brazil [8] to fit the transmission rates of malaria between mosquitoes and humans. Seasonality is incorporated through the mosquito biting rate in the fast subsystem so as to fit the seasonality of malaria cases in Brazil. We established a fit to the monthly incidence for Duffy-positive and Duffy-negative people, and determined a mean average percentage error of 7.02%. The transmission rates predicted by the parameter estimation are lower than the ones determined by literature. This leads to conservative ℛ_0_ values.

We investigate the impact of the fraction of people of XO and OO people on the reproduction number ℛ_0_. Fig. 8 are formulated by utilizing transmission rates obtained through parameter estimation, and identical transmission rates multiplied by 2.5 to yield a high transmission scenario. The left plot suggests for the transmission rates found through parameter estimation that the fraction of homozygous or heterozygous Duffy-positive people doesn’t affect ℛ_0_. The right plot with higher transmission rates illustrates that at least 85% of the population needs to be OO for ℛ_0_ *<* 1 or in other words *P. vivax* is controlled.

**Figure 8.**
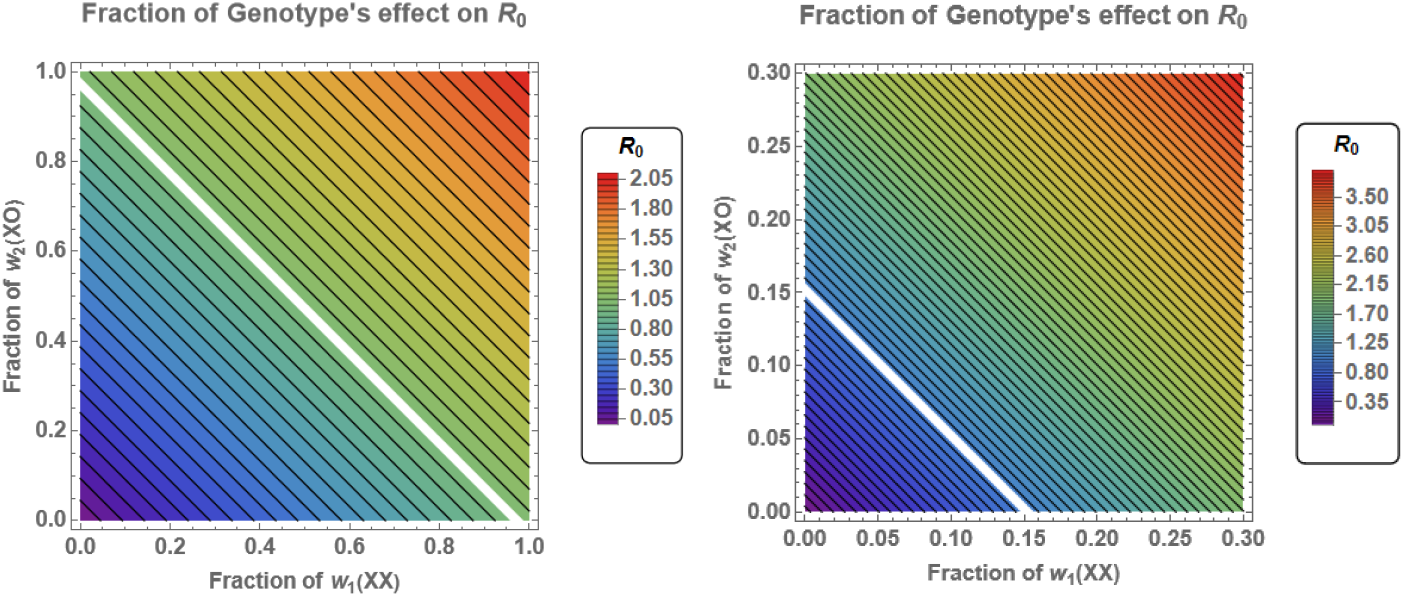
The white line through both figures is when ℛ_0_ = 1. The left figure is a contour plot of how the fraction of homozygous or heterozygous Duffy-positive people affects ℛ_0_ for low transmission rates. The right figure is the same but for higher transmission rates.

To quantify the relationship between disease burden and the frequencies of each Duffy genotype, we obtained Duffy genotype distributions for different locations from a handful of studies. In Fig. 9 is the map and the Duffy genotype breakdown of each sampled place. It can be seen that in the cities sampled from Cameroon and Kenya that the whole population is Duffy-negative while in the cities from China and Sweden the whole population is homozygous Duffy-positive. Some cities in Brazil have more diverse profiles such as Mazagão and Macapa.

**Figure 9.**
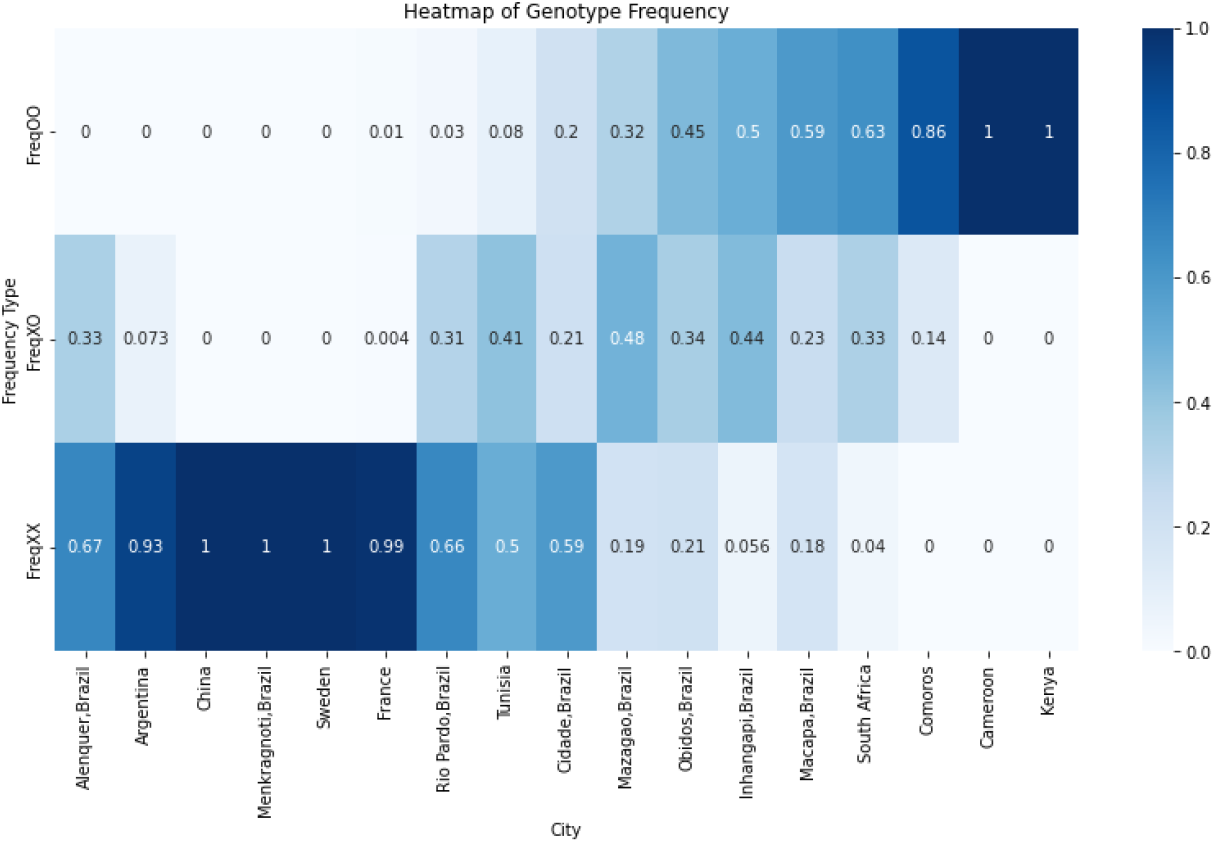
Frequencies of Duffy genotypes in different locations. Data from [13].

We then use ℛ_0_ defined in (8) to plot the basic reproduction number vs. the fraction of people who are Duffy-negative. Additionally, we use the model parameters from Table 3 and *w*_1_, *w*_2_, *w*_3_ from the data mentioned above. Note that some locations present in Fig. 9 have the same Duffy genotype frequencies as other locations, and are thus redundant and are not included in the scatterplot. To represent scenarios with high malaria transmission, we multiply the infection rate parameters *β*_*hi*_, *β*_*vi*_, (*i* ∈ {1, 2, 3 }) by a factor of 2.5 to increase the rates of infection.

In Fig. 10, the lines of best fit for the low transmission and high transmission scatterplots are *y* = − 0.8207*x* + 1.0644 and *y* = − 2.052*x* + 2.661, respectively. According to these slopes, a decrease of about 10 percentage points in the percentage of Duffy-negative people results in an increase of ℛ_0_ by about 0.08 when the malaria transmission is low. When we increase the transmission rates by a factor of 2.5, the same 10 percentage point difference causes a roughly 2.5 times larger increase to ℛ_0_ of about 0.21. Additionally, the relation is slightly nonlinear, as changes to smaller fractions of Duffy-negative people result in a smaller increase to ℛ_0_.

**Figure 10.**
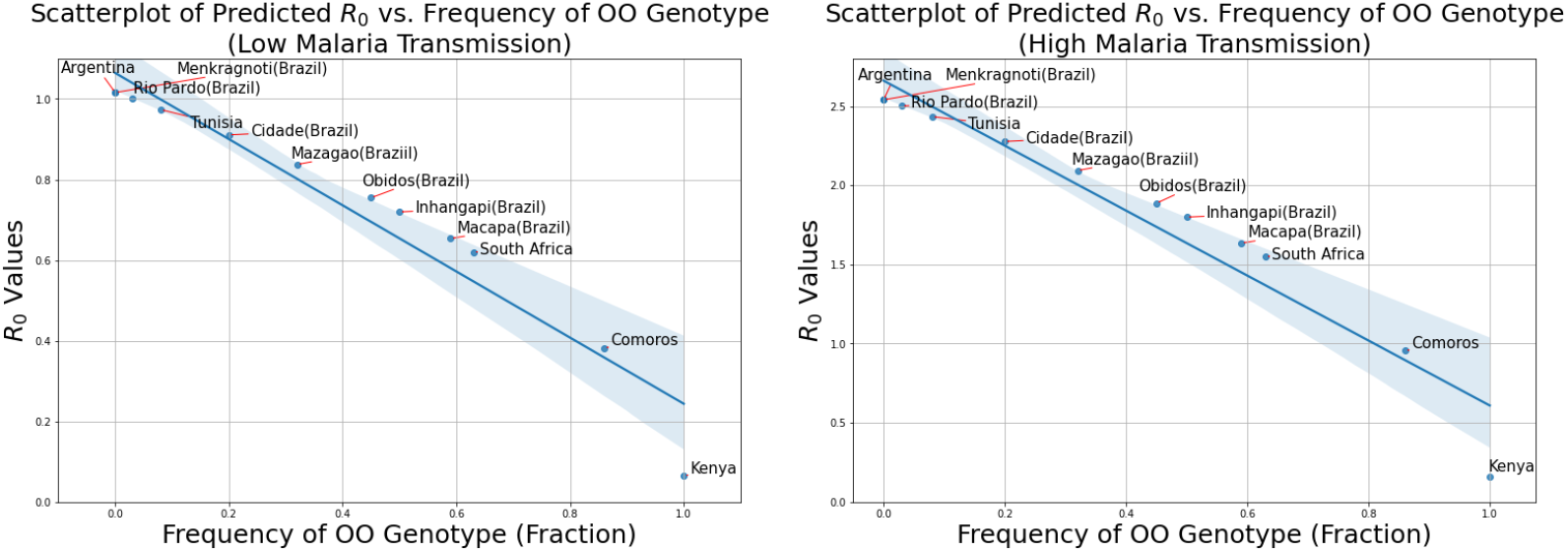
Scatterplots of predicted ℛ_0_ values versus the frequency of the OO genotype for low (Left) and high (Right) transmission rates with lines of best fit.

## 6 Discussion

The purpose of our mathematical model is to characterize the epidemiology of *P. vivax* malaria in relation to the DARC genotype. We do so by separating the genotype into three classifications, each relating to the number of alleles that are silent or expressed in a Duffy polymorphism. Thus, we employ a three-genotype model similar to the one Feng and Castillo-Chavez [5] developed to describe the sickle cell trait.

Using the slow system of our model, we obtained the fitness of the X-gene, which represents the likeliness of the expressed Duffy alleles FY*A and FY*B to persist into future generations. As shown in Fig. 4, as the recovery rate of Duffy-negative people (*γ*_3_) increases, it becomes more advantageous to be Duffy-negative and the X-gene is less likely to be selected for than the silent O-gene. As the infection rate of Duffy-negative people decreases, the fitness of the X-gene also decreases since it’s more beneficial to be Duffy-negative.

Through sensitivity analysis on both ℛ_0_ and the equation for the total number of infected people, we determined that the mosquito biting rate is highly influential parameter in both quantities, and thus highly influential in *P. vivax* malaria dynamics.

When we investigated the impact of the frequencies of the DARC genotypes and the transmission rates on ℛ_0_, we found that as the fractions of homozygous and heterozygous Duffy-positive people increase, ℛ_0_ also increases. Based on the model, there is an nonlinear inverse relationship between the frequency of Duffy negative people and the basic reproductive number ℛ_0_. To gather these results, we used a handful of Duffy genotype population breakdowns from various locations. Additionally, increasing the transmission rates resulted in an increase to the smallest percentage of Duffy-negative people required to keep *P. vivax* outbreaks under control.

Our model quantifies how the diversity of DARC genotypes impacts the dynamics of *P. vivax* malaria infections. In this paper, we established a framework to determine what percentage of Duffy-negative people are required to keep *P. vivax* outbreaks under control, once given the transmission rates from a specific geographic area. With statistically representative genetic data with respect to Duffy genotypes, our model can more accurately determine which populations are at risk of *vivax* malaria. Furthermore, once given region-specific infection parameters, this model can be applied to geographic regions around the world with varying genotypic frequencies. This provides valuable insight for the delegation of resources to different populations, such as antimalarial drugs, insecticide-treated bed nets, and malarial vaccine trials. Future work includes conducting Markov chain Monte Carlo sampling to establish confidence intervals for our parameter estimation. In addition, we hope to apply Extended Fourier Analysis Test (eFAST) to the fast subsystem with biting rate seasonality to refine the sensitivity analysis. Additionally, we will incorporate evolutionary game theory into our study, as it presents an interesting approach to further investigate the evolutionary dynamics of *P. vivax* malaria. Different seasonality models such as the two peak model outlined by White et al. [25] need to be considered as well.

## 7 Appendix

## 7.1 Mosquito Model Re-scale (3)

Taking the derivative of both sides with respect to time, substituting the expression for 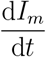 from (3), and simplifying yields

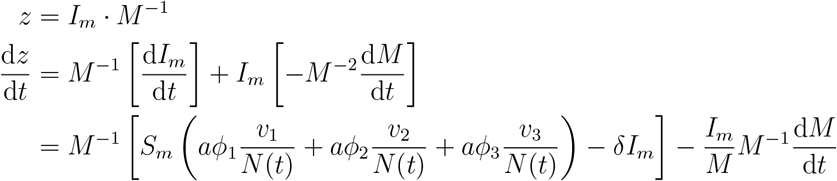

Note that *z* can be substituted for 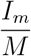 and (1 − *z*) for 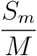 because of how we defined *z* and because *M* = *S*_*m*_ + *I*_*m*_.

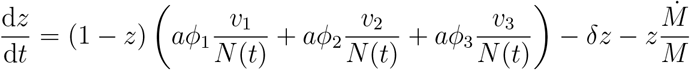

where 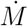 represents 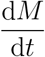. To maintain a roughly constant mosquito-to-human ratio (parameter *c*), we assume that *M* is constant. Implementing this assumption yields

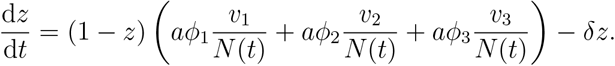

As such, our model can be expressed in the following equations.

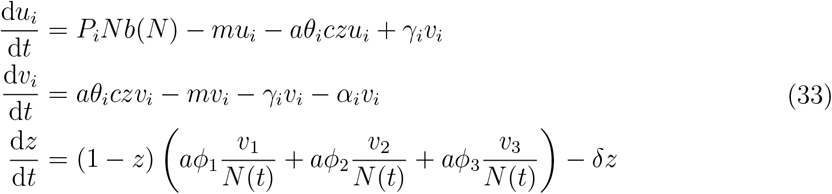

A similar process can be used to derive the equations for 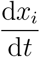 and 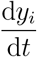 starting with the relations 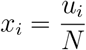 and 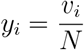. This results in the following fractional expressions of (3).

## Notes

### Competing Interest Statement

The authors have declared no competing interest.

